# Investigation of skin microbiota reveals *Mycobacterium ulcerans*-*Aspergillus* sp. trans-kingdom communication

**DOI:** 10.1101/869636

**Authors:** N. Hammoudi, C Cassagne, M. Million, S Ranque, O. Kabore, M. Drancourt, D. Zingue, A. Bouam

## Abstract

**Background:** *Mycobacterium ulcerans* secrete a series of non-ribosomal-encoded toxins known as mycolactones that are responsible for causing a disabling ulceration of the skin and subcutaneous tissues named Buruli ulcer. The disease is the sole non-contagion among the three most common mycobacterial diseases in humans. Direct contact with contaminated wetlands is a risk factor for Buruli ulcer, responsible for *M. ulcerans* skin carriage before transcutaneous inoculation with this opportunistic pathogen.

**Methodology and principal findings:** In this study, we analysed the bacterial and fungal skin microbiota in individuals exposed to *M. ulcerans* in Burkina Faso. We showed that *M. ulcerans*-specific DNA sequences were detected on the unbreached skin of 6/52 (11.5%) asymptomatic farmers living in Sindou versus 0/52 (0%) of those living in the non-endemic region of Tenkodogo. Then, we cultured the skin microbiota of asymptomatic *M. ulcerans* carriers and negative control individuals, all living in the region of Sindou. A total of 84 different bacterial and fungal species were isolated, 21 from *M. ulcerans*-negative skin samples, 31 from *M. ulcerans*-positive samples and 32 from both. More specifically, Actinobacteria, *Aspergillus niger* and *Aspergillus flavus* were significantly associated with *M. ulcerans* skin carriage. We further observed that *in vitro*, mycolactones induced spore germination of *A. flavus*, attracting the fungal network.

**Conclusion:** These unprecedented observations suggest that interactions with fungi may modulate the outcome of *M. ulcerans* skin carriage, opening new venues to the understanding of Buruli ulcer pathology, prophylaxis and treatment of this still neglected tropical infection.

**Author summary:** Buruli ulcer is a chronic infectious disease caused by the environmental opportunistic pathogen *Mycobacterium ulcerans* which secretes an exotoxin responsible for its pathogenicity. The reservoir and sources of *M. ulcerans* in the environment remain elusive and its mode of transmission is unclear. To acquire *M. ulcerans* infection, at least two conditions must be met, viable bacteria and a skin lesion as demonstrated by experimental animal models. In this study, we showed that *M. ulcerans* specific DNA sequences could be detected on the healthy skin of asymptomatic farmers living in one region of Burkina Faso where Buruli ulcer cases had already been reported, but not in Buruli ulcer-free regions, suggesting skin carriage after contacts with environmental sources. We also investigated the skin microbiota of *M. ulcerans* carriers and found significant associations of some bacteria and fungi with skin carriage of *M. ulcerans.* These associations may due to the effect of mycolactones on some fungi species. As we showed previously with *Mucor circinelloides* and here with *Aspergillus flavus*.

## Introduction

Buruli ulcer is a disabling chronic disease of the skin and the subcutaneous tissues [1]. The disease has been reported since 1948 in several tropical regions [2], with the highest incidence being observed in West African countries, including Côte d’Ivoire, Ghana, and Benin [3]. The causative agent *Mycobacterium ulcerans* is a non-tuberculous mycobacterium harbouring a 174-kb circular virulence plasmid pMUM [4-5] encoding three genes, *mls*A1, *mls*A2 and *mls*B, responsible for the non-ribosomal synthesis of mycolactone toxins, the main virulence factors of the pathogen [6]. This macrolide exotoxin is secreted by a group of closely related non-tuberculous mycobacteria named Mycolactone Producing Mycobacteria (MPM) [7-8]. Comparative analysis of whole genome sequences showed that MPM form a single clonal group that evolved from a *Mycobacterium marinum* parent [9]. This group is divided into three lineages, including frog and fish pathogens in one lineage [10-11], the Japanese strain *M. ulcerans* subsp. *shinshuense* in a second lineage, while the third lineage includes a highly clonal group responsible for Buruli ulcer in Africa and Australia [9]. Every strain synthesizes at least one of the eight congeners of mycolactone A/B named mycolactone C, D, E, F, dia-F, S1 and S2 [12]. Each type of mycolactone exhibits a variable degree of cytotoxicity, with mycolactone A/B being the most biologically active [13]. Indeed, this macrolide toxin is responsible for cytotoxic effects, namely, apoptosis and necrosis, in addition to immunosuppressive and analgesic effects [12] after *M. ulcerans* has penetrated breached skin to initiate discrete oedematous lesions that can evolve into typical Buruli ulcer lesions [14]. Currently, the mode of transmission of *M. ulcerans* is debated. While it is clear that *M. ulcerans* must be inoculated through the skin to elicit Buruli ulcer, the specific role of aquatic bugs and mosquitoes as potential vectors is debated along with the possibility of passive entry by any skin breach regardless of its cause. It is not known whether *M. ulcerans* can colonize skin in asymptomatic populations exposed to environments contaminated with *M. ulcerans*. Mycobacteria of the *M. ulcerans* group are known as environmental organisms residing in poorly defined aquatic ecological niches where they could be part of an alimentary chain [15]. We previously reported that *M. ulcerans* could thrive in environmental niches containing bacteria, fungi, algae and mollusks with which *M. ulcerans* may exchange nutrients [16]. One may hypothesize that the cutaneous microbiota, which is part of the individual, partially controls the expression of the *M. ulcerans* infection [17]. Indeed, bacteria and fungi present on skin contaminated with *M. ulcerans* could interact with the pathogen, as previously shown with the antagonism between *Staphylococcus lugdunensis* and *Staphylococcus aureus* in the nasal mucosa [18–19].

To contribute to this medical debate, we have undertaken the first study to evaluate the possibility of asymptomatic carriage of *M. ulcerans* on healthy skin, to characterize the cutaneous bacterial and fungal microbiota associated with the asymptomatic carriage of *M. ulcerans* and to assess the biological interactions between *M. ulcerans* and the skin microbiota.

## Methods

### Ethics Statement

This study was approved by the Centre MURAZ Ethics Committee, Burkina Faso, and reference 2018-11/MS/SG/CM/CEI.

All adult subjects provided informed oral consent, and a parent or guardian of any child participant provided informed oral consent on the child’s behalf after explaining the merits of the study. Written consent could not be obtained because the study was conducted among an illiterate population. The research presents no more than minimal risk of harm to subjects and involves no procedures for which written consent is normally required outside of the research context (Sec. 56.109 IRB review of research).

#### Sample collection

Sampling was performed in two regions of Burkina Faso. Sindou is located in the rural district of Niofila, Douna Department, Province of Léraba in the Cascades region in the southwest of Burkina Faso near the borders with Côte d’Ivoire and Mali. People sampled in this region are farmers in frequent contact with stagnant water because of their daily activities in rice, banana and vegetable cultivation areas that are irrigated from a nearby dam. Cases of Buruli ulcer have already been reported in this region [20]. Tenkodogo, a town located in the province of Boulgou and the Central-Eastern region of Burkina Faso, was used as a negative control region free of Buruli ulcer. Additionally, in this region, farmers working near two dams in rice and vegetable cultures were swabbed (the lower legs of the farmers were cleaned with water and then swabbed with sterile swabs (Deltalab, Barcelona, Spain) containing 1 mL of sterile 0.9% sodium chloride solution, the swabbed parts did not contain any visible lesions.). A total of 104 farmers were sampled, 52 farmers in each region. In Sindou region, the average age was 37 years, 30 females and 22 males. In Tenkodogo region the male/female ratio was 27/25 respectively with an average age of 20 years. One sample was taken per individual.

### Detection of *M. ulcerans* DNA

#### Real-time PCR amplifications

Total DNA from the samples and from a six-week-old culture of *M. ulcerans* CU001 (positive control) was extracted using a QIAamp Tissue Kit by QUIAGEN-BioRobot EZ1, according to the manufacturer’s instructions (Qiagen, Hilden, Germany). To assess PCR inhibition, 10 µL of an external control was added to 190 µL of sample volume as previously described [21. Extracted DNA was used in real-time PCR (RT-PCR) to amplify the insertion sequences (IS*2404* and IS*2606*) and the ketoreductase-B domain of the mycolactone polyketide synthase genes (KR-B) [22] using RT-PCR reagents from Roche PCR Kit (Roche Diagnostics, Meylan, France) and primers and probes as previously described [21] in a CFX 96™ real-time PCR thermocycler and detection system (BIO-Rad, Marnes-la-Coquette, France). To estimate *M. ulcerans* inoculum in skin swabs, we performed three calibration curves for the IS*2404*, IS*2606*, and KR-B genes. The total DNA of the *M. ulcerans* CU001 strain calibrated at 1 McFarland = 10^6^ CFU was extracted with EZ (Qiagen, Hilden, Germany). Then, 10-fold serial dilutions of up to 10^−8^ were made to generate a calibration curve for each system. Two reactional mixes were incorporated into each PCR run as negative controls. Samples were considered positive when the KR-B gene was detected with Ct < 40 cycles, and at least one of the two PCRs of the insertion sequence gave Ct < 40 cycles and KR-B was detected with Ct < 40 cycles.

### Skin microbiota repertoire

#### Bacterial culture

All RT-PCR-positive samples collected in Sindou were selected for culture. Six negative samples collected in the same region were used as negative controls. For each sample, a cascade dilution (up to 10^−10^) was performed in sterile PBS (Thermo Fisher Diagnostics, Dardilly, France), and 100 µL of each dilution was inoculated in duplicate on blood agar plates (bioMérieux, Marcy l’Etoile, France). One blood agar plate was incubated aerobically, and the second was anaerobically incubated at 37°C for 72 hours. After incubation, each colony presenting a unique morphology was sub-cultured onto a blood agar plate and incubated at 37°C for 48 hours under the appropriate atmosphere to isolate each colony type.

#### Fungal culture

*M. ulcerans* PCR-positive samples were cultured in three different culture media, namely, homemade RMI medium, Sabouraud agar (Oxoid, Dardilly, France) and Chromagar (Becton Dickinson, Le Pont de Claix, France) incubated at 30°C for seven days. Each colony was sub-cultured and incubated under the same conditions. The six PCR-negative samples (negative controls) were treated using the same protocol.

#### Matrix assisted laser desorption ionization time-of-flight identification of colonies

The identification of the bacterial and fungal colonies was carried out using matrix-assisted laser desorption ionization time-of-flight mass spectrometry MALDI-TOF-MS as previously described [23,24]. For bacteria, each colony was deposited in duplicate onto a MALDI-TOF MSP 96 target plate (Bruker Daltonics, Leipzig, Germany), and 2 μL of matrix solution (saturated solution of alpha-cyano-4-hydroxycinnamic acid in 50% acetonitrile and 2.5% trifluoroacetic acid) was added to each spot and allowed to dry for 5 mins and then analysed by Microflex spectrometer (Bruker Daltonics) using the software MALDI BioTyper 3.0 (Bruker Daltonics). For the identification of fungi, each colony was incubated in 1 mL of 70% ethanol for 10 min and then centrifuged at 1,300 g for 5 min. The pellet was treated with 20 μL of acetonitrile and formic acid (v.v) at 100% and 70%, respectively. This mixture was then centrifuged at 1300 g for 5 min, and 1.5 μL of the supernatant was deposited on a MALDI-TOF-MS target and allowed to dry before 1.5 μL of matrix was deposited on each spot, allowed to dry for 5 min and then analysed by par Microflex spectrometer (Bruker Daltonics) using the software MALDI BioTyper 3.0 (Bruker Daltonics).

#### Molecular identification and sequencing

All colonies that remained unidentified by MALDI-TOF-MS were subjected to molecular identification by sequencing the bacterial 16S rRNA [25] and the fungal ITS1, ITS2, β tubulin and TEF regions. The primers used in this study are reported in Table S1. DNA extraction was performed using BioRobot EZ1 (Qiagen, Les Ullis, France) using the commercial EZ1 DNA Tissue Kit according to the manufacturer’s instructions (Qiagen). PCR was performed using Hotstar *Taq* polymerase according to the manufacturer’s instructions (Qiagen) using a thermocycler (Applied Biosystem, Paris, France). PCR products were separated by electrophoresis on a 1.5% agarose gel and stained with SYBR® safe (Thermo Fisher Scientific) before being visualized under an ultraviolet transilluminator. PCR products were then purified using a Millipore NucleoFast 96 PCR kit following the manufacturer’s recommendations (Macherey-Nagel, Düren, Germany) and sequenced using the BigDye Terminator Cycle Sequencing Kit (Applied Biosystems) with an automatic sequencer ABI (Applied Biosystems). Sequences were assembled using the software ChromasPro 1.7 (Technelysium Pty Ltd., Tewantin, Australia) and blasted in the NCBI databank (http://blast.ncbi.nlm.nih.gov/Blast.cgi) to identify bacterial species and against the Mycobank database http://www.mycobank.org/ to identify fungal species.

#### Fungi*-M. ulcerans* biological interactions

The three fungal species found to significantly correlate with the presence / absence of *M. ulcerans* skin carriage (*A. flavus*, *A. niger* and *P. rubens*) were cultured in the presence / absence of mycolactones AB / C extracted from a culture of *M. ulcerans* CU001 to observe the effect of mycolactones on spore germination and the fungal network. Briefly, agar plates were cut into T-shape, the fungal spores were placed on the middle strand, and two virgin absorbent paper discs were placed on each side of the plates. 20uL of PBS were put on the first disc (negative control) and 20uL of mycolactones AB / C on the second disc. The chemoattractant effect of mycolactone AB / C was measured as previously described [25]. On the other hand, fungal spores were put in Sabouraud liquid medium containing 20uL of mycolactones AB / C or 20uL of PBS as a negative control. The effect of Mycolactone AB/C on spore germination was then monitored as previously described [26].

### Statistical analyses

The first exploratory step of unsupervised analysis of the cutaneous bacterial and fungal microbiota used a main component analysis by integrating the presence or absence of each of the fungi detected in at least one individual and adding the variable "detection of *M. ulcerans* by PCR". We complemented this analysis with a supervised comparison between individuals positive and negative for *M. ulcerans* by PCR. To do this, we compared the detection frequency in the two groups. Statistical significance was calculated by the two-sided exact Fisher’s test.

Owing to the possible risk of missing important findings, adjustments for multiple comparisons were not performed, as suggested for exploratory work [27]. A double-clustering heatmap was used to visualize the potential clustering of cultured fungal and bacterial repertoires with *M. ulcerans*-positive or negative skin samples. Finally, we carried out a discriminant factor analysis to test whether the fungal skin microbiota discriminated against individuals who were positive or negative for *M. ulcerans* skin carriage. The principal component and discriminant factor analyses were performed using XLSTAT v2019.1 statistical and data analysis solution (Long Island, NY, USA (https://www.xlstat.com)). Multivariate analysis was performed via the Genmod procedure with the SAS 9.4 statistical software using a negative binomial distribution, adjusted for the effect of time (quantitative variable) and by using generalized estimating equations to account for the non-independence of repeated measures to analyse the results of germination assays. The Wilcoxon signed rank test and Chi-squared test were used to compare the attraction effect of fungi by mycolactones and to compare *M. ulcerans* prevalence in the two geographical regions of Sindou and Tenkodogo, respectively. Statistical tests were performed on the website biostaTGV, https://biostatgv.sentiweb.fr.

## Results

### *M. ulcerans* DNA detection

Negative controls introduced in every batch of real-time PCR remained negative, and positive controls were positive for all the targeted sequences IS*2404*, IS*2606* and KR-B, allowing the establishment of calibration curves. Of the 52 swabs collected in the region of Sindou, eight were positive for IS*2404*, six were positive for IS*2606*, two were positive for both IS*2404* and IS*2606*, one was positive for both IS*2404* and KR-B, two were positive for both IS*2606* and KR-B, one was positive for all three targeted sequences and 32 samples were negative (Fig. 1). Therefore, 6/52 (11.5%) swabs were positive for at least two *M. ulcerans* DNA sequences. As expected, only three samples were positive for IS*2404* only in Tenkodogo, so none of the 52 skin samples collected in this negative control region were positive for *M. ulcerans*, confirming the probable absence of *M. ulcerans* in this region. The difference in the prevalence of RT-PCR-based detection of *M. ulcerans* between the two regions was significant (P = 0.04, N-1 Chi-squared test). Based on the calibration curves established during this study, the Ct values observed here were extrapolated to 9-50 colony forming units (CFU) of *M. ulcerans*.

**Fig. 1:**
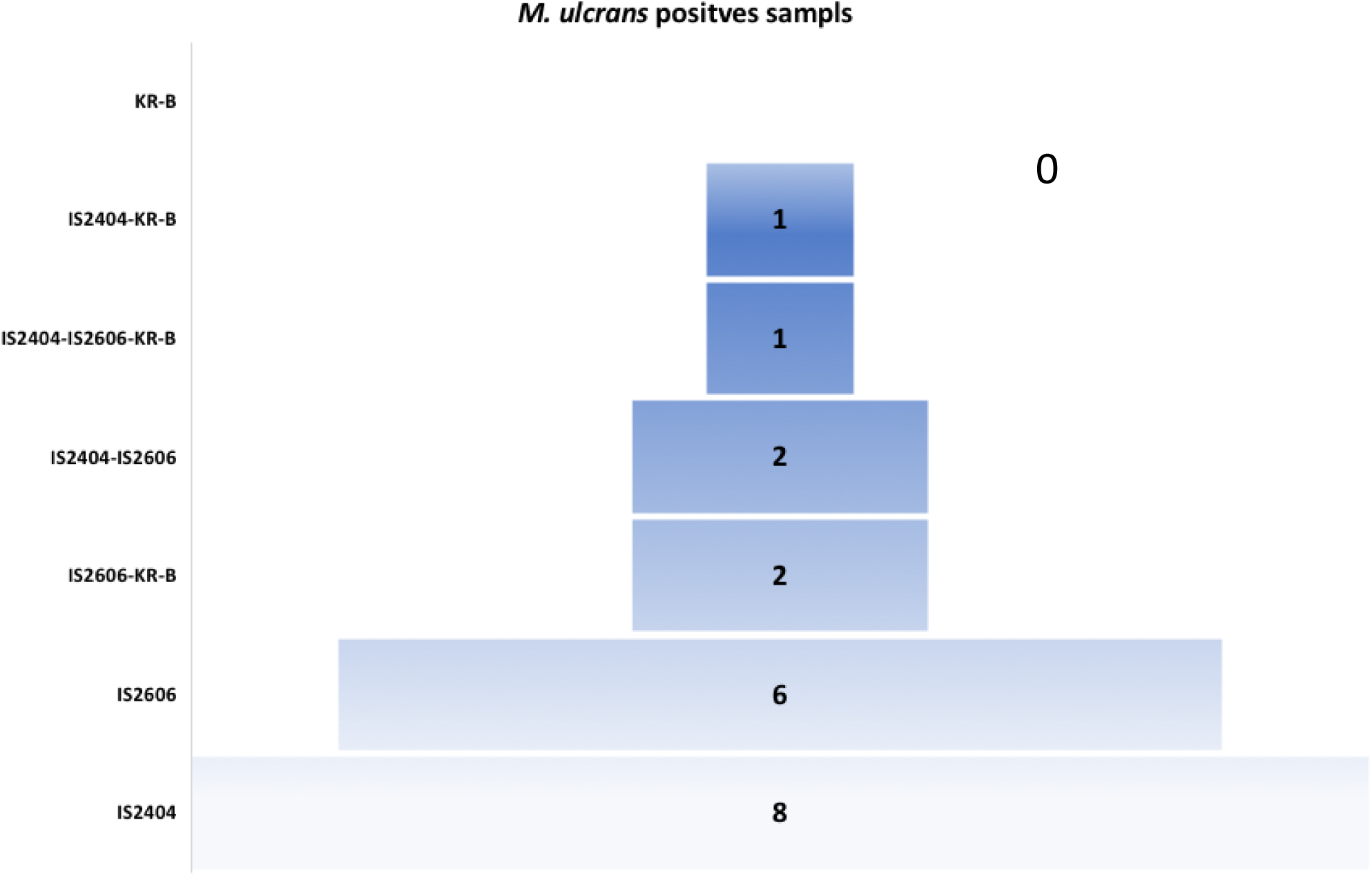
Real-time PCR detection of *M. ulcerans* from 52 swabs collected in the Sindou region, IS*2404* and IS*2606* are specific insertion sequences for *M. ulcerans* KR-B ketoreductase-B gene.

### Skin microbiota identification

A total of 84 different species of microorganisms (62 bacterial and 22 fungal species) belonging to 45 different genera were recovered from the twelve samples. Thirty-one different species of microorganisms (19 bacterial and 12 fungal species) grew exclusively in *M. ulcerans*-PCR positive samples. Thirty-two different species (22 bacterial and 10 fungal species) grew only from *M. ulcerans*-PCR negative samples. Twenty-one bacterial species grew on both *M. ulcerans*-PCR positive samples and *M. ulcerans*-PCR negative samples. No fungal species were found in both types of samples simultaneously (Fig. 2). Globally, the microbiota of *M. ulcerans*-PCR positive samples was statistically enriched for Actinobacteria (Fisher’s test p-value = 0.03). The genus *Aspergillus* and the species *Aspergillus flavus* were significantly associated with *M. ulcerans-*PCR positive samples (two-sided Fisher’s exact test p-value = 0.0021 and 0.015, respectively), and the genus *Penicillium* was significantly associated with *M. ulcerans*-PCR negative samples (p-value = 0.0021, Fisher’s exact test). All *Zygomycetes, Acidovorax, Brevundimonas, Cutibacterium*, and *Homoserinibacter* species were recovered from PCR *M. ulcerans*-positive samples, whereas all *Penicillium, Cellulosimicrobium, Franconibacter, Ochrobactrum, Porphyromonas, Roseomonas, Achromobacter* and *Lelliottia* species were recovered from *M. ulcerans*-PCR-negative samples, but these associations were not statistically significant (Fig. 2). The dendrogram obtained by the agglomerative hierarchical classification of the species recovered from samples harboured two distinct clusters perfectly separating the *M. ulcerans*-PCR positive samples from the *M. ulcerans*-PCR negative samples (Fig. 3). These results highly suggest that the skin microbiota was significantly correlated with *M. ulcerans* carriage.

**Fig. 2:**
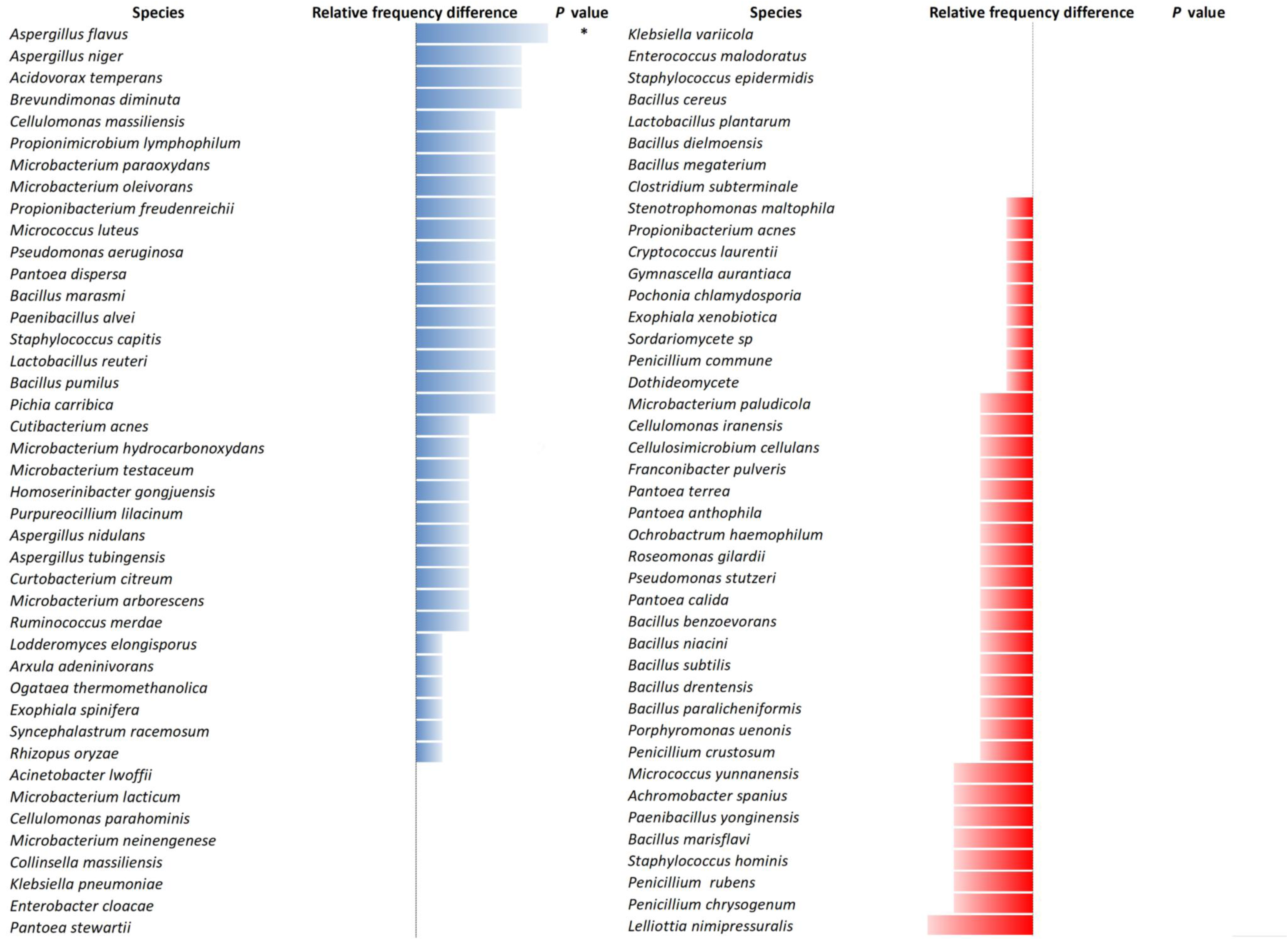
The different microorganisms grown from 12 skin samples comprising 6 *M. ulcerans* positive samples and 6 negative samples are presented with their relative frequency differences and associated P-values. A total of 84 species distributed among 62 bacteria and 22 fungi were cultured.

**Fig. 3:**
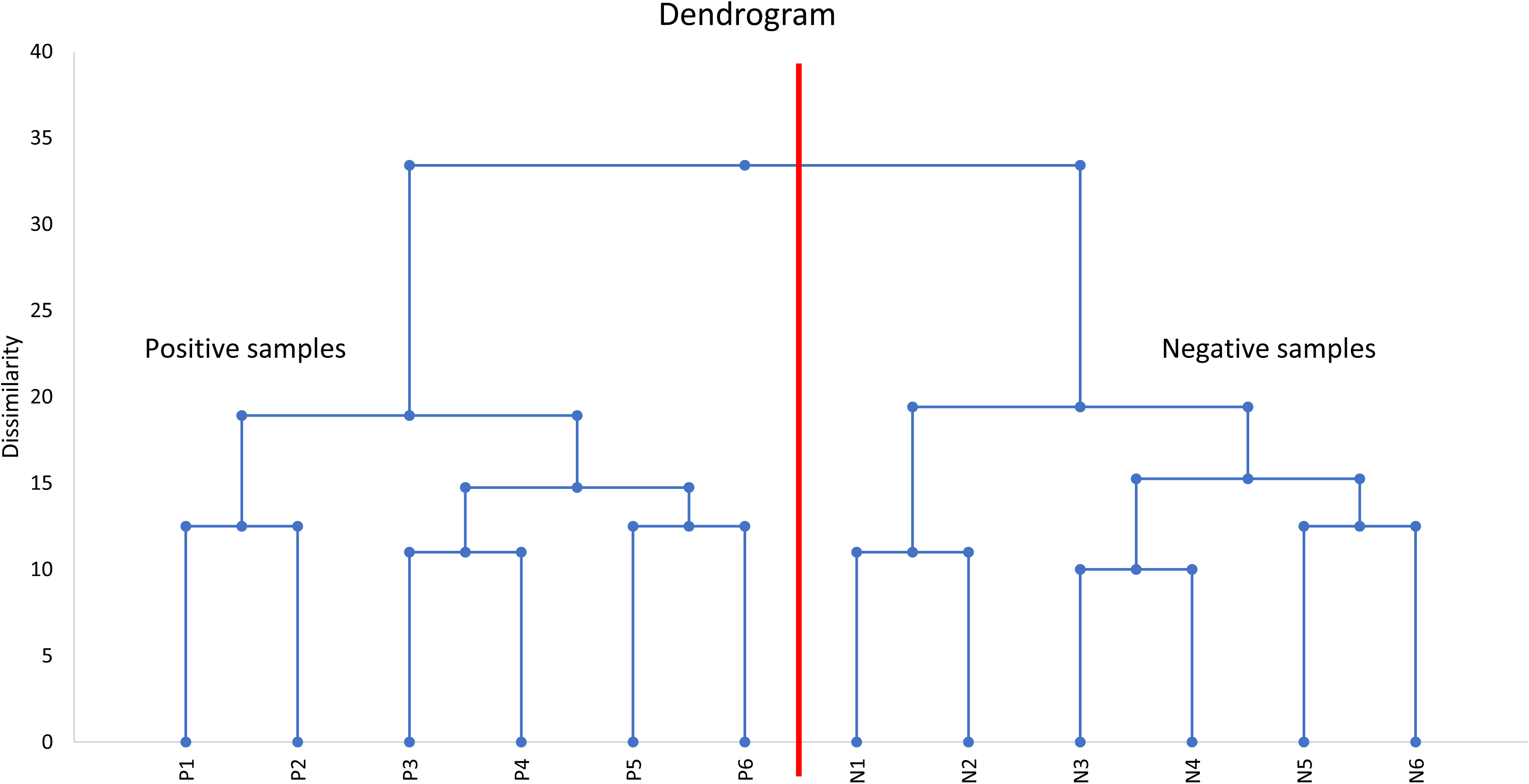
The agglomerative hierarchical classification dendrogram of individuals according to their cutaneous microbiota associated with the detection (P1 to P6) or absence (N1 to N6) of *M. ulcerans. M. ulcerans* PCR-negative samples and *M. ulcerans* PCR-positive samples were clearly separated into two distinct clusters according to the composition of the skin microbiota.

### Principal component analysis

After submission of the results of fungal and bacterial culturomics and PCR *M. ulcerans* to PCA analysis, two varifactors (F1 and F7; representing both a total of 28.3% of the variance of the data) were selected because these two varifactors represented the highest variance of the variable “PCR *M. ulcerans*”. F1 and F7 contributed 20.4% and 7.9% of the overall variability, respectively. *Aspergillus flavus*, *Propionimicrobium lymphophilum* and *Propionimicrobium freudenreichii* were strongly associated with a positive PCR for *M. ulcerans*, whereas *Penicillium rubens*, *Penicillium chrysogenum* and *Lelliota nimipressuralis* were associated with a negative PCR. Surprisingly, detection of *Aspergillus* fungi positively correlated with PCR detection of *M. ulcerans*. Indeed, *A. flavus* was the fungus exhibiting the strongest positive association with *M. ulcerans*. On the other hand, and unexpectedly, we observed that the detection of *Penicillium* fungi was anti-correlated with the detection of *M. ulcerans*. No *Penicillium* detection positively correlated with *M. ulcerans* detection, while *P. rubens* and *P. chrysogenum* were the only fungi to anti-correlate with *M. ulcerans* (Fig. 4).

**Fig. 4:**
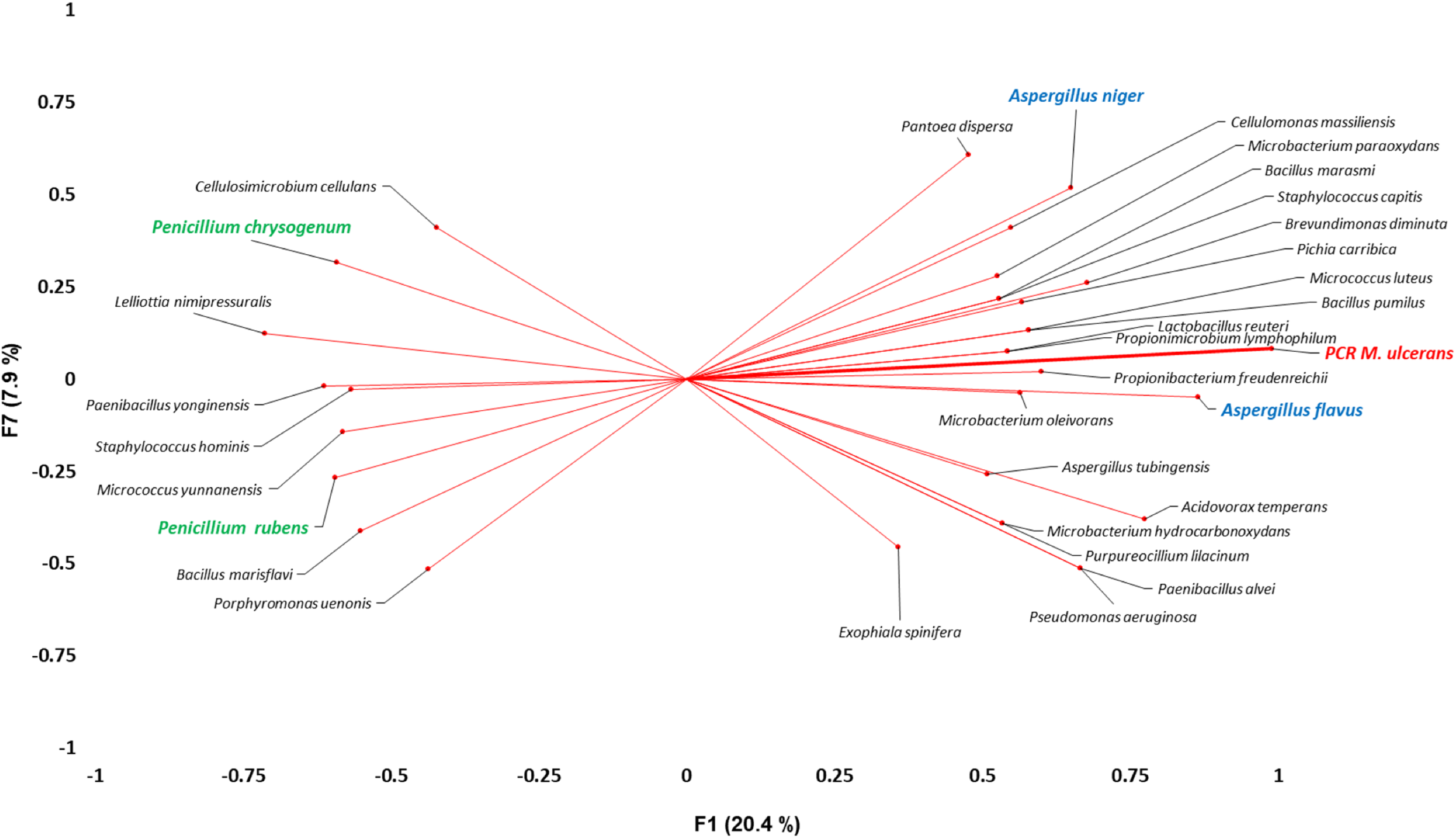
PCA of 84 bacterial and fungal species isolated from individuals in Burkina Faso revealed a strong association of *A. flavus*, *P. lymphophilum* and *P. freudenreichii* with *M. ulcerans* skin carriage.

### Factor discriminant analysis

Interestingly, factor discriminant analysis identified five fungal and bacterial species that significantly discriminated *M. ulcerans*-PCR positive and negative samples, including *A. flavus, Brevundimonas diminuta, Propionimicrobium lymphophilum, Pantoea dispersa* and *P. rubens* (p-values: 0.001, 0.01, 0,049, 0.049, 0.049, respectively –Table S2). The discriminant analysis showed that the cutaneous fungal and bacterial microbiota discriminated with 100% accuracy between *M. ulcerans* positive and negative groups (receiver operator curve analysis: area under curve = 1 - no error in the confusion matrix – perfect clustering).

### Fungi*-M. ulcerans* biological interactions

Mycolactones significantly increased spore germination of A. *flavus* after a 10-hour incubation (stimulated 91.2% vs. control 28.75%; P <.0001), significantly increased spore germination of *A. niger* (stimulated 75.42% vs. control 69.17%; P <.0001), and significantly decreased spore germination of *P. rubens* (stimulated 45.2% vs. control 50.7%; P <.0001) (Fig. 5). Moreover, we observed that mycolactones significantly attracted *A. flavus* and *A. niger* during the T-test assay (*P*-value: 0.013 and 0.003, respectively) (Fig. 6).

**Fig. 5:**
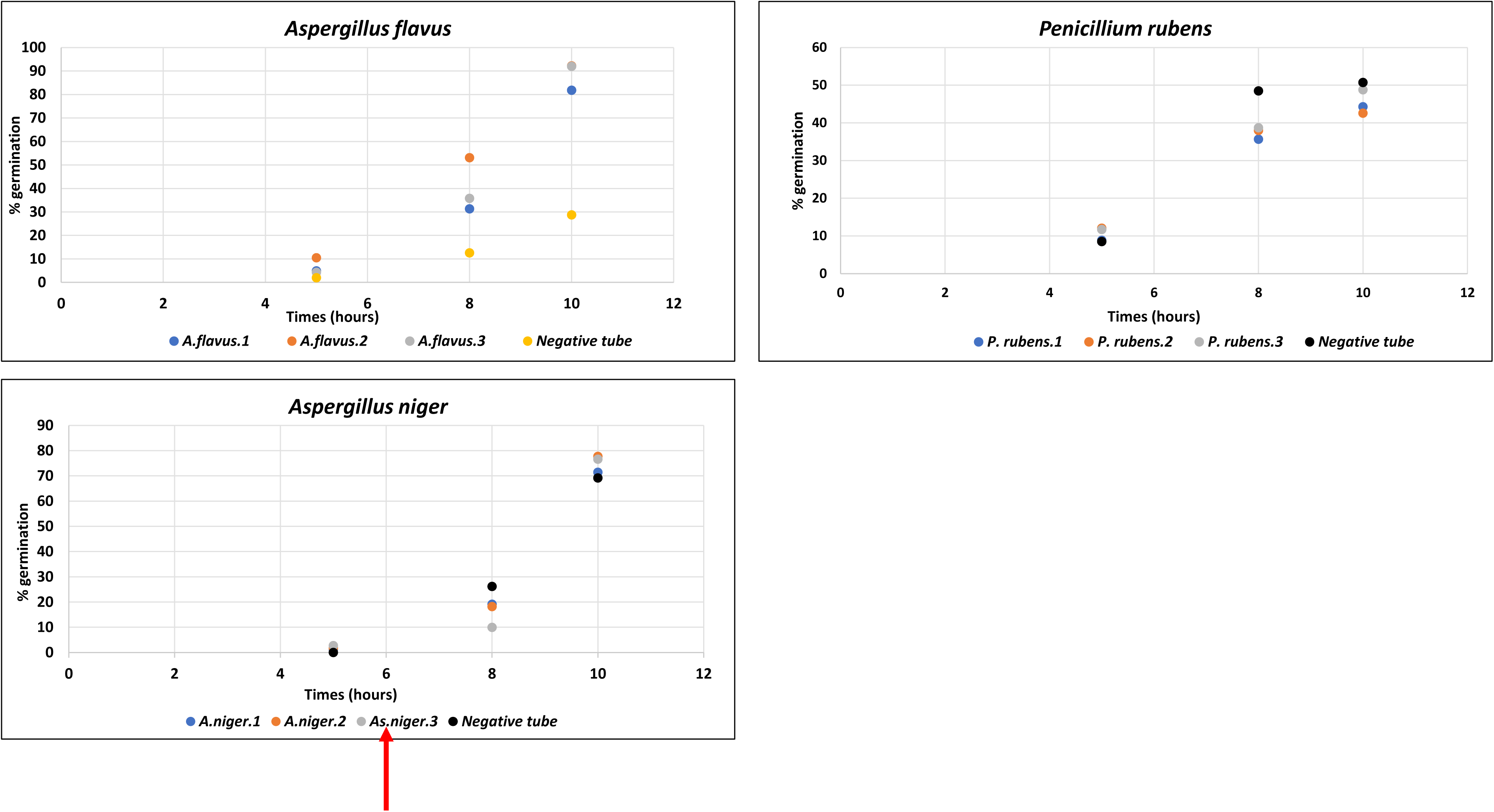
Mycolactone-induced spore germination of *A. flavus*, *A niger* and *P. rubens* in the presence of PBS as a negative control.

**Fig. 6:**
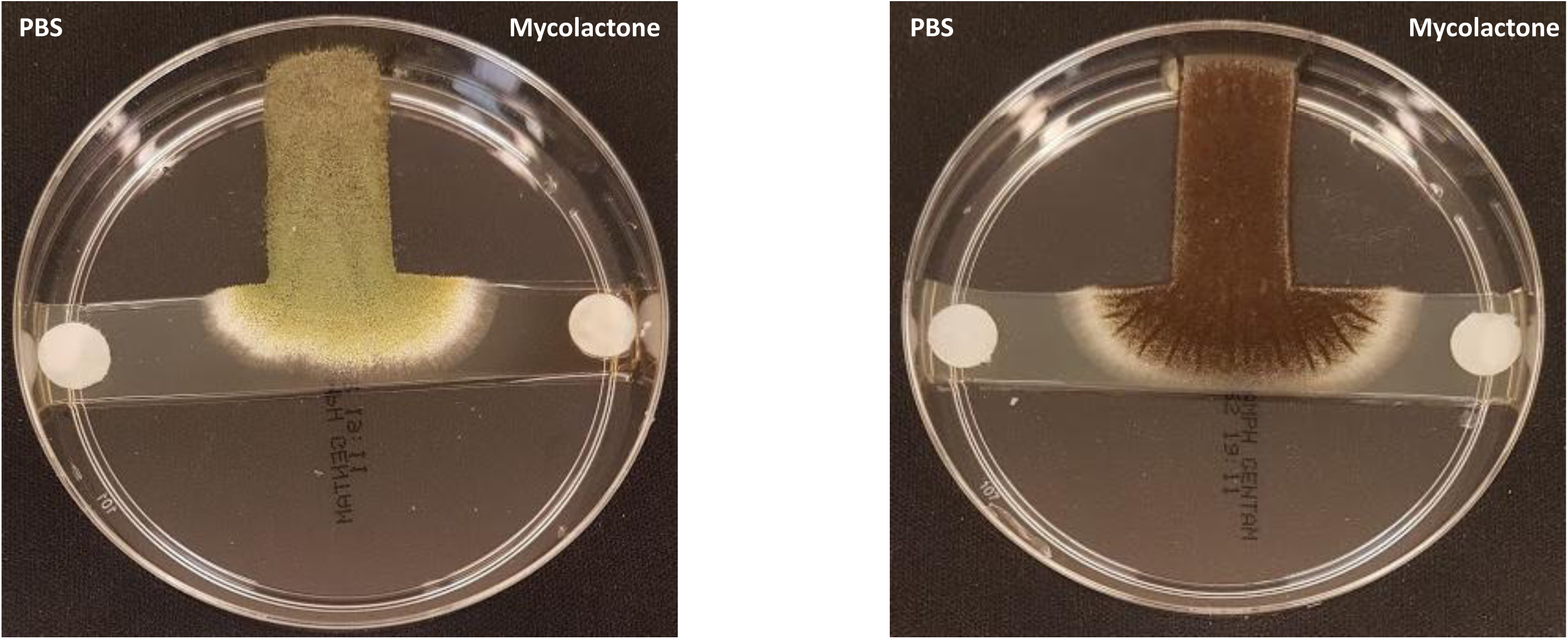
Fungi attraction test: mycolactones (right side disk) attract *A. flavus* and *A. niger* in the presence of PBS as a negative control (left side disk).

## DISCUSSION

Using a standard RT-PCR assay to detect the asymptomatic skin carriage of *M. ulcerans* in skin samples collected from individuals living in Burkina Faso, we detected a prevalence of 11.5% of *M. ulcerans* carriage in individuals residing in the Sindou region, Burkina Faso, where Buruli ulcer cases have been previously reported [20]. Our data agree with a previous report of cutaneous carriage of *M. ulcerans* in Ghana among people practising agriculture without any protective clothing and in infants crawling on the soil [28]. Thus, the data reported here confirm that in Buruli ulcer-endemic areas, some people are asymptomatic skin carriers of *M. ulcerans*. The *M. ulcerans* inoculum we detected on the skin has been reported to be sufficient to initiate Buruli ulcer lesions in a murine model [29]. These observations suggest that asymptomatic skin carriage could be a previously undescribed condition in the natural history of Buruli ulcer.

Exploration of the skin microbiota at the interface of healthy and diseased skin is in its infancy, especially regarding Buruli ulcer [30]. Comparison of skin microbiota between *M. ulcerans*-PCR positive and negative samples revealed a specific cutaneous microbiota associated with asymptomatic *M. ulcerans* skin carriage, even predicting *M. ulcerans* skin carriage. Other skin pathogens and diseases (leprosy and psoriasis) have been previously related to a specific skin microbiota [31-32]. The novelty in our study was to find significant, antiparallel associations between *M. ulcerans* and fungi: *M. ulcerans* was detected on the skin along with *Aspergillus*, in which spore germination and the fungal network were stimulated by mycolactones, whereas the detection of *M. ulcerans* has never been associated with the the presence of *Penicillium* species in the skin, Accordingly, spore germination of *P. rubens* was inhibited by mycolactones. Therefore, our clinical observations were not fortuitous but revealed transkingdom mycobacteria-fungi interactions, supporting preliminary observations made with the Zygomycete *Mucor circinelloides* [26]. Of note, the transkingdom communication concerned *P. rubens*, a penicillin-producing strain, alias *P. chrysogenum*, made famous by Sir Alexandre Fleming [33].

All these observations suggest for the first time that transkingdom communications between fungi and mycobacteria are of medical interest, partially driving the natural history of Buruli ulcer. These observations will stimulate additional studies to disclose whether this holds true for other mycobacterial infections of medical interest.

## ACKNOWLEDGEMENTS

The authors acknowledge the technical help of Nicholas Armstrong.

This work was supported by the French Government under the « Investissements d’avenir » (Investments for the Future) programme managed by the Agence Nationale de la Recherche (ANR, fr: National Agency for Research), (reference: Méditerranée Infection 10-IAHU-03). This work was supported by Région Sud (Provence Alpes Côte d’Azur) and European funding FEDER PRIMMI.

## Supplementary data

**Supplementary Table 1:**
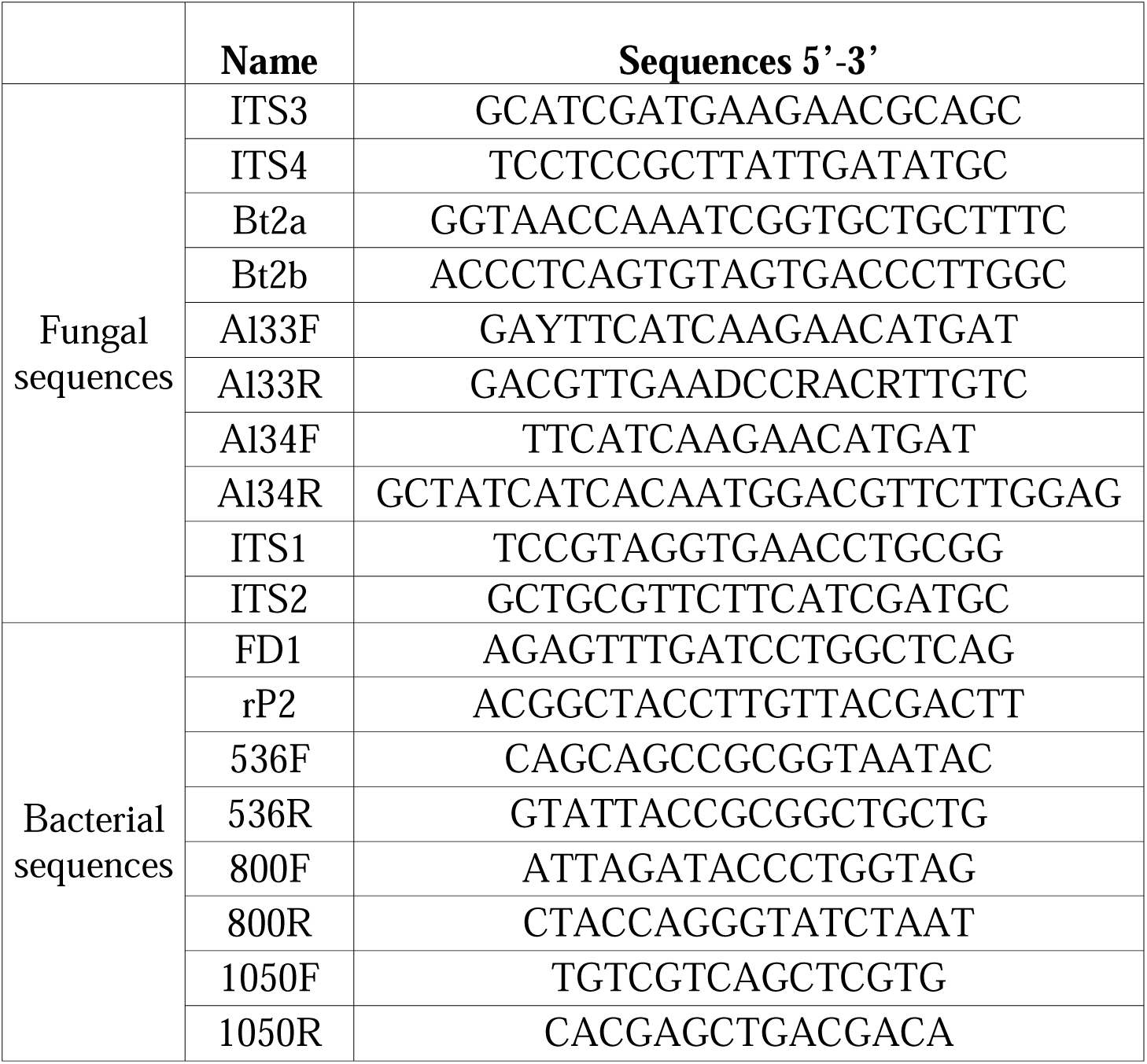
Primers used in this study for the PCR-based identification of fungi and bacteria.

**Supplementary Table 2:** Factor discriminant analysis (FDA) revealed five fungal and bacterial species that significantly discriminated *M. ulcerans*-PCR positive and negative samples.

| Variable | Lambda | F | DDL1 | DDL2 | p-value |
| --- | --- | --- | --- | --- | --- |
| *Propionimicrobium lymphophilum*-0 | 0,667 | 5,000 | 1 | 10 | 0,049 |
| *Brevundimonas diminuta*-0 | 0,500 | 10,000 | 1 | 10 | 0,010 |
| *Pantoea dispersa*-0 | 0,667 | 5,000 | 1 | 10 | 0,049 |
| *Penicillium rubens*-0 | 0,667 | 5,000 | 1 | 10 | 0,049 |
| *Aspergillus flavus*-0 | 0,286 | 25,000 | 1 | 10 | 0,001 |

